# Is single-step genomic REML with the algorithm for proven and young more computationally efficient when less generations of data are present?

**DOI:** 10.1101/2022.01.19.476983

**Authors:** Vinícius Silva Junqueira, Daniela Lourenco, Yutaka Masuda, Fernando Flores Cardoso, Paulo Sávio Lopes, Fabyano Fonseca e Silva, Ignacy Misztal

## Abstract

Efficient computing techniques allow the estimation of variance components for virtually any traditional dataset. When genomic information is available, variance components can be estimated using genomic REML (GREML). If only a portion of the animals have genotypes, single-step GREML (ssGREML) is the method of choice. The genomic relationship matrix (**G**) used in both cases is dense, limiting computations depending on the number of genotyped animals. The algorithm for proven and young (APY) can be used to create a sparse inverse of **G**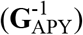 with close to linear memory and computing requirements. In ssGREML, the inverse of the realized relationship matrix (**H**^-1^) also includes the inverse of the pedigree relationship matrix, which can be dense with long pedigree, but sparser with short. The main purpose of this study was to investigate whether costs of ssGREML can be reduced using APY with truncated pedigree and phenotypes. We also investigated the impact of truncation on variance components estimation when different numbers of core animals are used in APY. Simulations included 150K animals from 10 generations, with selection. Phenotypes (h^2^ = 0.3) were available for all animals in generations 1-9. A total of 30K animals in generations 8 and 9, and 15K validation animals in generation 10 were genotyped for 52,890 SNP. Average information REML and ssGREML with **G**^-1^ and 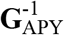 using 1K, 5K, 9K, and 14K core animals were compared. Variance components are impacted when the core group in APY represents the number of eigenvalues explaining a small fraction of the total variation in **G**. The most time-consuming operation was the inversion, with more than 50% of the total time. Next, numerical factorization consumed nearly 30% of the total computing time. On average, a 7% decrease in the computing time for ordering was observed by removing each generation of data. APY can be successfully applied to create the inverse of the genomic relationship matrix used in ssGREML for estimating variance components. To ensure reliable variance component estimation, it is important to use a core size that corresponds to the number of largest eigenvalues explaining around 98% of total variation in **G**. When APY is used, pedigrees can be truncated to increase the sparsity of **H** and slightly reduce computing time for ordering and symbolic factorization, with no impact on the estimates.

**Lay Summary:** The estimation of variance components is computationally expensive under large-scale genetic evaluations due to several inversions of the coefficient matrix. Variance components are used as parameters for estimating breeding values in mixed model equations (MME). However, resulting breeding values are not Best Linear Unbiased Predictions (BLUP) unless the variance components approach the true parameters. The increasing availability of genomic data requires the development of new methods for improving the efficiency of variance component estimations. Therefore, this study aimed to reduce the costs of single-step genomic REML (ssGREML) with the Algorithm for Proven and Young (APY) for estimating variance components with truncated pedigree and phenotypes. In addition, we investigated the influence of truncation on variance components and genetic parameter estimates. Under APY, the size of the core group influences the similarity of breeding values and their reliability compared to the full genomic matrix. In this study, we found that to ensure reliable variance component estimation it is required to consider a core size that corresponds to the number of largest eigenvalues explaining around 98% of the total variation in **G** to avoid biased parameters. In terms of costs, the use of APY slightly decreased the time for ordering and symbolic factorization with no impact on estimations.

**Teaser Text:** Estimation of variance components is becoming computationally challenging due to the increasing size of genomic information. We investigated the impacts of using the algorithm for proven and young (APY) in genetic evaluations. The use of APY has no impact on variance components and genetic parameters estimation.

## Introduction

Restricted maximum likelihood (REML), described by Patterson and Thompson (1971), is a popular method for parameter estimation. Because it uses the mixed model equations (Henderson, 1975), it is resistant to selection bias, and efficient implementations are currently available. With the Average Information (AI) algorithm, convergence is often achieved in a few rounds. With traces obtained by sparse matrix factorization and inversion (Meyer, 1997), computing variance components is feasible even with large models.

When genomic information is available, two versions of REML may be applicable. When only genotyped animals have phenotypes, genomic REML (GREML) can be applied with a genomic relationship matrix (**G**). In general, such a matrix is dense, and the cost of dense matrix operations would limit computations depending on the models. When only a fraction of animals are genotyped, a single-step genomic REML is applicable (ssGREML). In the latter, the combined relationship matrix (**H**) has dense blocks due to the genomic information, limiting the efficiency of sparse matrix operations. Lately, Masuda et al. (2015) developed a sparse matrix package YAMS that identifies dense blocks and computes them efficiently. For ssGREML, with genomic computation, such a package resulted in up to 100 times speedup, allowing four trait models with 20,000 genotyped animals (Masuda et al., 2015).

In general, it is of interest to include many genotyped animals in parameter estimation and evaluations to account for genomic selection or pre-selection (Patry and Ducrocq, 2011). For instance, the greatest reliability in a single-step genomic BLUP was obtained using 50% of the heritability computed with a non-genomic REML (Misztal et al., 2017). The number of genotyped animals is increasing fast for some species. As an example, almost 3 million Holsteins have been genotyped in the US (https://queries.uscdcb.com/Genotype/cur_freq.html). However, the cost of dense matrix operations with **G** in REML using YAMS is quadratic for memory and cubic for operations, which limits computations to around 50,000 animals.

The genomic information has a limited dimensionality due to the limited effective population size (Stam, 1980; VanRaden, 2008; Misztal, 2016). Such dimensionality varied from 4,000 in pigs and chickens to 15,000 in Holsteins (Pocrnic et al., 2016c). Assuming limited dimensionality, the inverse of **G** (**G**^-1^) – as needed by REML – can be sparsely constructed using the APY algorithm, with close to linear memory and computing requirements. Subsequently, the inverses for over 2 million animals can be computed and stored (Tsuruta et al., 2021). However, the inverse of **H** also includes the inverse of a pedigree-based relationship matrix for genotyped animals (Aguilar et al., 2010). Such a matrix can be dense with a long pedigree, but it is sparser with a shorter pedigree. Thus, it could not be efficiently stored in large populations but had to be accommodated indirectly (Strandén and Mäntysaari, 2014; Masuda et al., 2017).

The first purpose of this study was to find whether the costs of ssGREML can be reduced using the APY algorithm with truncated pedigree and phenotypes. We hypothesize the truncation could help to preserve the system’s sparsity, given that APY **G**^-1^ is sparser than the inverse of the pedigree relationship matrices for deep pedigrees. The second purpose was to investigate to what extent such truncation influences variance components and heritability estimates when different numbers of core animals are used in APY.

## Material and Methods

Animal care and use committee approval was not needed because data were simulated.

### Data simulation

To evaluate the computational effectiveness of the proposed approach for estimating variance components using genomic information, we simulated data using the QMSim software (Sargolzaei et al., 2011). The simulator generated a historical population undergoing drift and mutation and a recent population undergoing selection. The historical population consisted of 1,000 generations with a constant size of 50,000 individuals. Then, 800 more generations were simulated where the number of individuals was reduced to 20,000, mimicking a bottleneck event. The recent population (P1) consisted of 20 males and 15,000 females randomly sampled from the last historical generation based on high phenotypic values. Individuals were mated along ten generations producing a litter size of 1 with an equal probability of being male or female, following a random mating design. Moreover, we considered a sire replacement rate of 0.50 and a dam replacement rate of 0.20. Genomic information was available for 45,000 animals from generations 8 through 10 (three youngest generations).

A total of 29 chromosomes of different lengths (ranging from 40 to 146 cM) were simulated. Biallelic markers (n = 52,890) were evenly spaced along the chromosomes with equal frequency in the first generation of the historical population. Potentially, 1,242 quantitative trait loci (QTL) affected the trait and explained all the additive genetic variation; QTL allele effects were sampled from a Gamma distribution with a shape parameter of 0.4. The mutation rate for markers (recurrent mutation) and QTL was assumed to be equal to 2.5 × 10^−5^ per locus per generation (Solberg et al., 2008).

The simulated trait had phenotypic variance and mean of 1.0, heritability and QTL heritability of 0.30, and residual variance of 0.70. The simulated phenotypes were composed of:

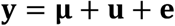

where **y** is the vector of phenotypes, **μ** is the vector of overall mean, **u** is the vector of weighted sum of QTL effects (i.e., additive genetic effect or animal effect), **e** is the vector of residuals.

The standard error of estimates was small using 5 replicates during preliminary investigations of this study. Because of that, the results are based on one replicate.

### Variance components

Variance components were estimated using the average information (AI) REML algorithm as implemented in the AIREMLF90 software (Misztal et al., 2002), which was modified to incorporate the YAMS package (Masuda et al., 2014; Masuda et al., 2015). The incorporation of YAMS was essential for this kind of task when using genomic information. The package applies the supernodal method using multi-core optimized libraries (i.e., parallel computing). The most computationally expensive part of the variance component estimation is obtaining the inverse of the coefficient matrix used in traces. To that, efficient algorithms are used to invert large and sparse matrices, which are based on three steps *(i)* ordering, *(ii)* factorization (i.e., symbolic and numerical), and *(iii)* sparse inversion. Ordering is not mandatory, but it saves a large amount of memory and time in the factorization step as it reduces the *fill-in* effect (zero elements in the original matrix could become nonzero elements in the factorized matrix). This effect can be minimized by ordering using appropriate techniques. In the next step, the coefficient matrix (LHS of the mixed model equations) is factorized into two triangular matrices by LU decomposition – L matrix. Finally, the Takahashi algorithm can be used for inversion. The supernodal method is expected to provide faster inversions because they find and process dense blocks in sparse matrices. Note that LHS inversion is only required to estimate variance components or compute prediction error variance (PEV, obtained from diagonal elements of an inverted LHS). If the objective is to solve the system of equations to obtain breeding values, iterative methods as the preconditioned conjugate gradient (Lidauer et al., 1999; Tsuruta et al., 2001) can be efficiently applied.

The model used to estimate variance components was based on the single-step method, in which the inverse of the realized relationship matrix (**H**^−1^) is used in the mixed model equations instead of **A**^−1^. Single-step genomic BLUP (ssGBLUP) is used for breeding value estimation, whereas ssGREML is used for variance components estimation. The inversion of **H** is computed as follows (Aguilar et al., 2010):

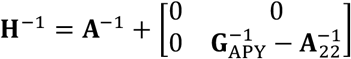

where **A**^−1^ is the inverse of the pedigree relationship matrix, 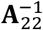 is the inverse of the pedigree relationship matrix for genotyped animals, computed by the algorithm described in Colleau (2002). The genomic relationship matrix (**G**) was computed as follows:

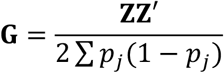

where **Z** is the matrix of gene content centered by the current allele frequencies, and *p*_*j*_ is the allele frequency of SNP *j*. Inbreeding coefficients were considered when constructing the three relationship matrices. This provides a better equivalence between genomic and pedigree-based relationship matrices, leading to a more similar genetic base (Aguilar et al., 2020). The 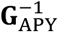 is the inverse of the genomic relationship matrix obtained using the algorithm for proven and young (APY) (Misztal et al., 2014; Misztal, 2016). This algorithm considers that genotyped individuals are arbitrarily divided into core (c) and noncore (*n*). Breeding values for noncore (**u**_*n*_) can be described as a linear function of breeding values of core (**u**_*c*_):

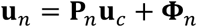

where **P**_*n*_ = **Z**_*n*_(**Z′**_*c*_ **Z**_*c*_ + **I***α*)^−**1**^**Z′**_*c*_ is a matrix that relates breeding values of noncore and core, and **Φ**_*n*_ is the mendelian sampling term which has non-diagonal variance but can be approximated to diagonal. In cases where the number of core is large enough, breeding values of noncore depend only on breeding values of core (see Misztal (2016) for additional details). The inverse of **G**_APY_ is constructed as following:

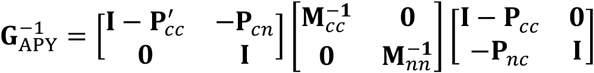

If 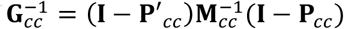is known, the complete inverse can be simplified to:

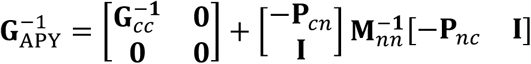

where 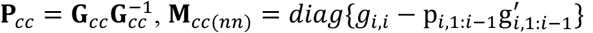 for individual *i* in the core (noncore) group. Because 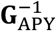 is conditioned only on the genotypic information of core animals, the matrix is sparser than the full **G**^−**1**^ regularly used in ssGBLUP (Misztal, 2016). Note that the covariance between two noncore individuals is null, but variances are stored in the matrix.

The construction of the genomic matrix using APY in BLUPF90 software can be done in two possible implementations. The first construction builds a single matrix for all core and noncore. The second construction builds the genomic matrix in blocks and it aims to save computing memory as it require less operations than single matrix (Masuda et al., 2016). Currently, the single matrix construction is implemented for variance component estimation.

### Scenarios

The scenarios below were built to evaluate the impact of the (1) size of the core group in APY and the (2) influence of skipping zero elements from the LHS under different amounts of pedigree and phenotypic data used in variance components estimation.

#### Core group of different sizes

Pocrnic et al. (2016a) evaluated the prediction accuracy using APY in simulation tests.

The authors suggested the greatest accuracy was found by selecting the number of core individuals equal to the number of largest eigenvalues explaining 98% of **G** (a number from now on referred to as eigen98). This study tested core groups of different sizes to evaluate the impact on variance components and heritability estimates. A total of four scenarios were tested by allocating 1K (one thousand), 5K, 9K, and 14K randomly sampled out of 45,000 genotyped individuals. For each of those scenarios, the largest variation explained was 72.03% (eigen70), 91.09% (eigen90), 95.70% (eigen95), and 98.07% (eigen98), respectively. For computational reasons, the singular value decomposition of **Z** was calculated instead of the eigenvalue decomposition of **G**.

#### Evaluating the influence of pedigrees and phenotypes

Using 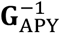 helps to reduce computing time for genomic predictions because of its sparsity (Fragomeni et al., 2015; Masuda et al., 2016); however, in the single-step approach, the combined **H**^−1^ contains also **A**^−1^ and 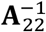, which are relatively dense. The APY method was earlier applied to the construction of 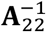 without success (Breno Fragomeni, personal communication). Although the sparsity of 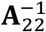 may not be a requirement for genomic predictions, it becomes essential for reducing computing time for variance components estimation to follow the sparsity of 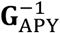. A reduction in the number of generations was attempted to increase the sparsity in **A**^−1^ and 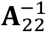 total of seven different scenarios were designed, differing on the number of pedigree generations used for variance components estimation. Reduction in the generations of phenotypes was also used to follow pedigree incompleteness and avoid bias. The scenarios were designed to mimic a real situation where the actual founder population is usually unknown. Only three genotyped generations (45,000 most recent animals) were kept in the genomic file for further analyses. Subsequent scenarios were constructed by removing one generation of phenotypes and pedigree at a time, from the oldest to the youngest animals.

#### The influence of zero elements in the Mixed Model Equations (MME)

Lastly, a scenario aimed to evaluate the impact of discarding zero elements from the LHS of MME on computing performance and variance components estimation. For that, the *OPTION skip_zero_in_dense_matrix* was used in AIREMLF90 (Misztal et al., 2014) to store only non-zero elements of 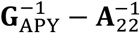. When this option was used, the scenario was termed “Reduced”, and otherwise “Full”.

## RESULTS AND DISCUSSION

Previous studies have investigated the properties of APY, including its implementation for large-scale genomic evaluations (Fragomeni et al., 2015; Lourenco et al., 2015; Masuda et al., 2016) and its efficiency in real and simulated populations with different effective population sizes (Pocrnic et al., 2016b; Pocrnic et al., 2016c). Bradford et al. (2017) studied the impact of different core definitions, and Misztal et al. (2020) evaluated the GEBV fluctuation when changing the core group in APY. Additionally, Vandenplas et al. (2018) investigated the impact of using APY on GEBV estimation in crossbreeding schemes; Hidalgo et al. (2021) compared the GEBV variation due to the inclusion of new data and changing the APY core animals. Finally, Lourenco et al. (2018) studied the impact of using 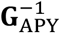 instead of **G**^−1^on the estimation of SNP effects. Our study evaluated the feasibility of using APY for variance components estimation, the impact of removing generations of pedigree and phenotypic data on computing time, and the influence of using a different number of core animals to construct the genomic matrix. Variance components were estimated using AIREML modified to incorporate the YAMS package for sparse matrix calculations (Masuda et al., 2014).

### Heritability estimates and computing performance

Heritability, residual variance, and additive variance estimated using a different number of generations in the pedigree and cores sizes in APY are shown in Figures 1-3. The standard deviation of variance components and heritability across generations is shown in Table 1. Because the simulation involved a certain level of selection, the expected heritability should slightly deviate from the simulated value of 0.3. Therefore, the scenario with 10 generations of data (i.e., full pedigree and full phenotypes) was used as a benchmark.

**Table 1.** Standard deviation of variance components and heritability calculated across generations using a complete (Full) mixed model equations (MME), and a reduced MME after skipping zero elements (Reduced).

| Parameter<sup>1</sup> | Core<sup>2</sup> | Scenario |  |
| --- | --- | --- | --- |
|  |  | Full | Reduced |
| $\sigma_a^2$ | 1K | 0.037 | 0.037 |
|  | 5K | 0.011 | 0.013 |
|  | 9K | 0.008 | 0.008 |
|  | 14K | 0.005 | 0.005 |
| $\sigma_e^2$ | 1K | 0.028 | 0.028 |
|  | 5K | 0.007 | 0.007 |
|  | 9K | 0.005 | 0.005 |
|  | 14K | 0.000 | 0.004 |
| $h^2$ | 1K | 0.032 | 0.032 |
|  | 5K | 0.011 | 0.011 |
|  | 9K | 0.005 | 0.005 |
|  | 14K | 0.005 | 0.005 |

<sup>1</sup> $\sigma_a^2$ : additive variance, $\sigma_e^2$ : residual variance, $h^2$ : heritability
<sup>2</sup> Number of individuals included as core to build the inverse of genomic matrix using the algorithm of proven and young (APY)

**Figure 1.**
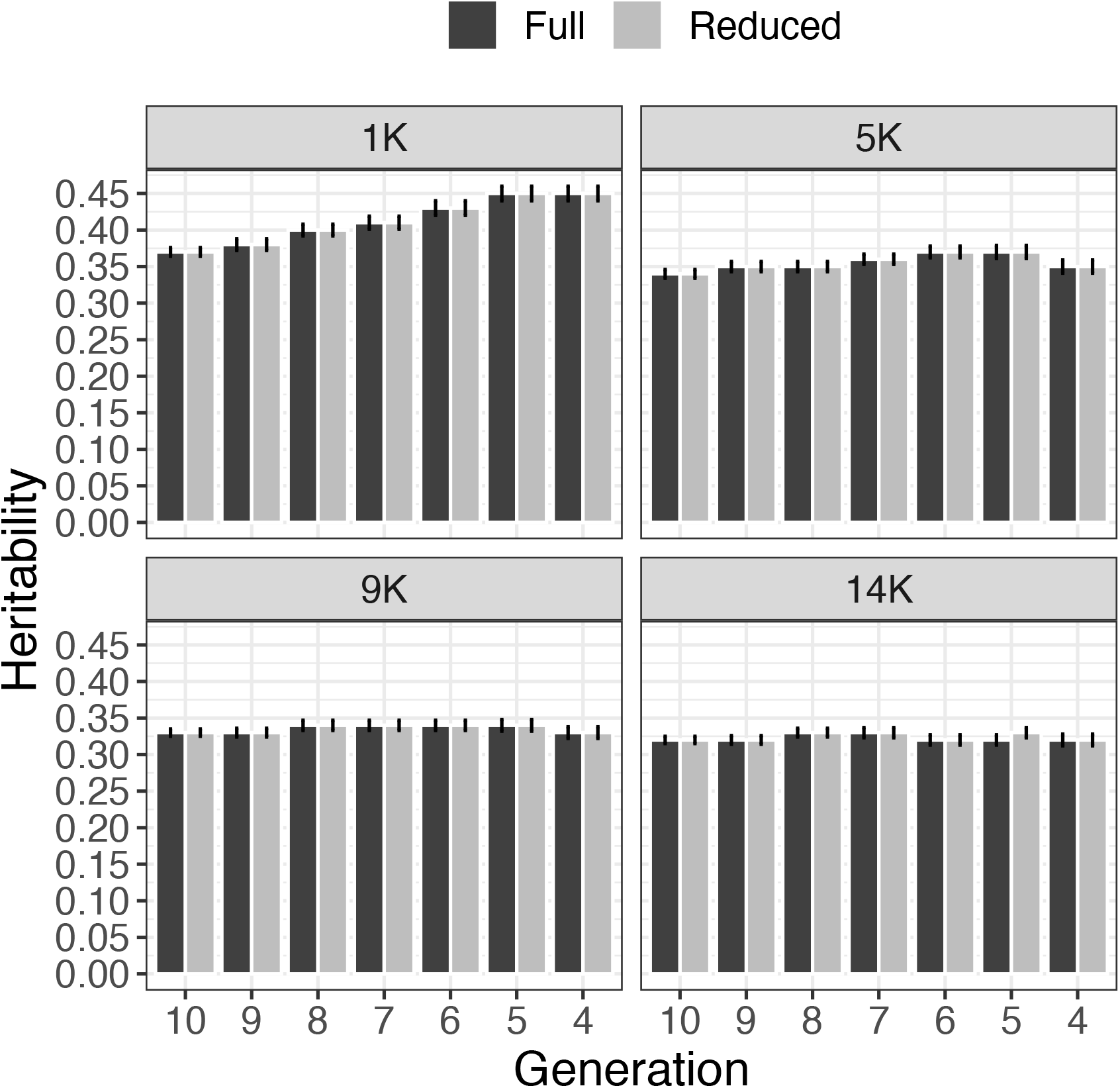
Heritability calculated along one replicate of simulation considering different number of generations with pedigree and phenotypic data under different number of core animal in APY. Two scenarios were considered, where zeros were stored (Full) or not (Reduced). Error bars represents the standard error of prediction under REML.

**Figure 2.**
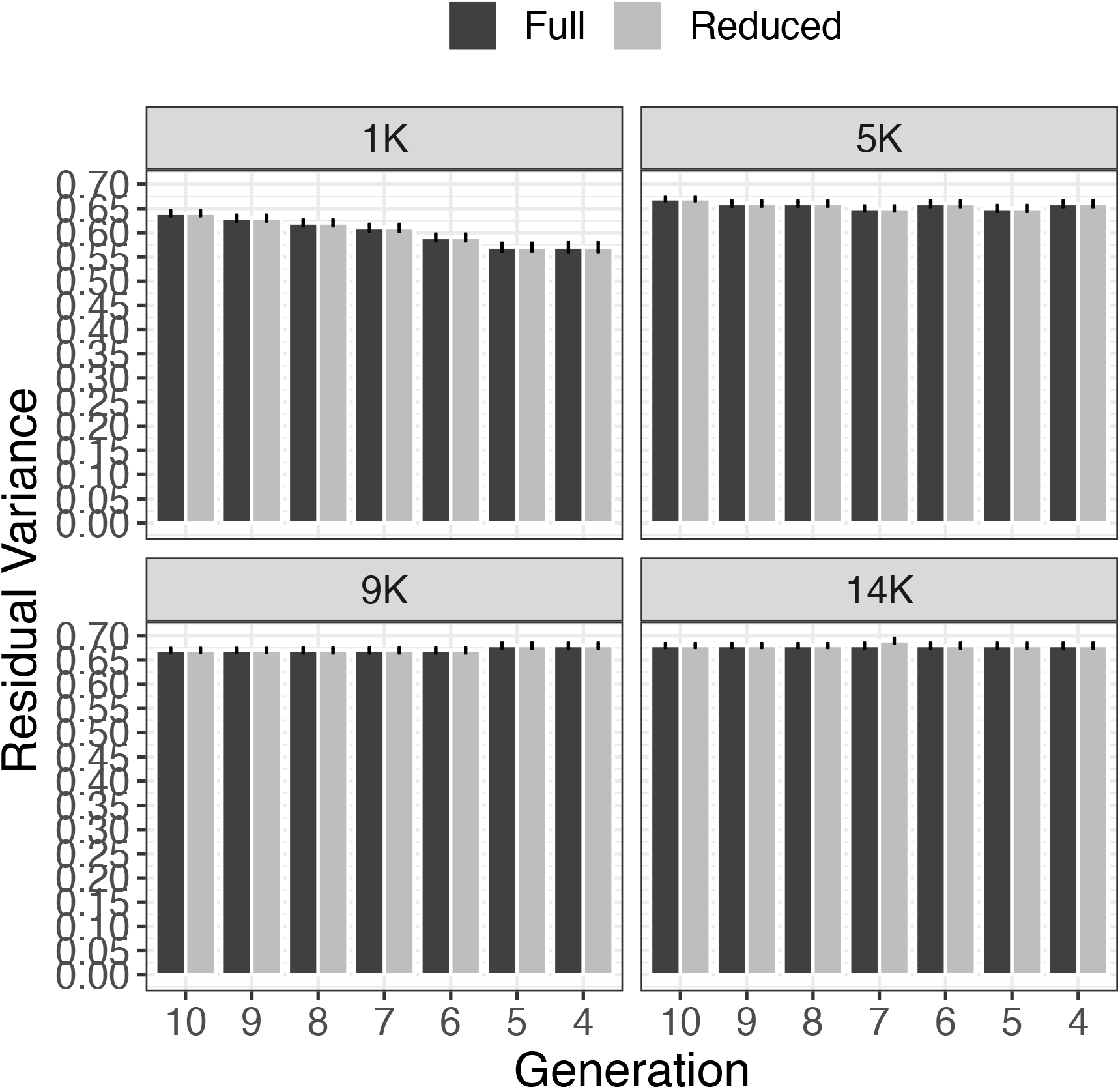
Residual variance calculated along one replicate of simulation considering different number of generations with pedigree and phenotypic data under different number of core animal for APY calculation. Two scenarios were considered, where zeros were stored (Full) or not (Reduced). Error bars represents the standard error of prediction under REML.

**Figure 3.**
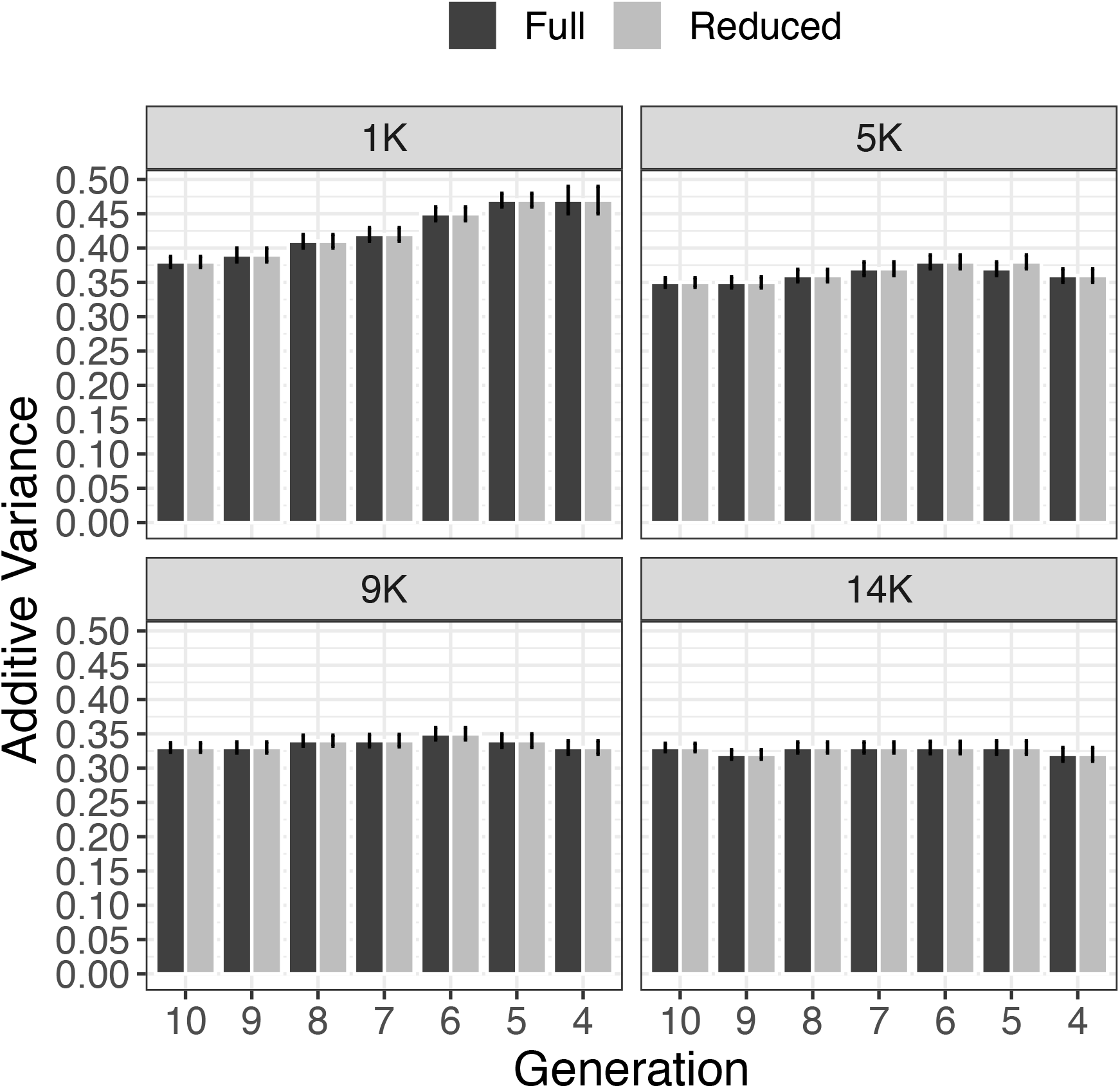
Additive variance calculated along one replicate of simulation considering different number of generations with pedigree and phenotypic data under different number of core animal in APY. Two scenarios were considered, where zeros were stored (Full) or not (Reduced). Error bars represents the standard error of prediction under REML.

In general, the variance components and heritability estimates approached the simulated values as the number of core approached eigen98. The scenario using 1K individuals (i.e., eigen70) in the core was the most sensitive to removing generations, suggesting that variance components are highly impacted when the core group in APY represents the number of eigenvalues explaining a smaller fraction of the total variation in **G**. From a prediction accuracy standpoint, a similar behavior was also observed in other studies (Pocrnic et al., 2016a; Pocrnic et al., 2016c); however, the impact on variance components had not been investigated before.

Although pedigrees were more limited after removing a few generations of data, the combination of pedigree and genomic information and the use of a proper core size controlled the bias in variance components and heritability estimation. Small fluctuations on variance components and heritability were observed when retaining only 4 to 6 generations of pedigree and phenotypes with a core size equal to eigen98. In these scenarios, the difference in heritability was almost nonexistent; this was also true when comparing Full and Reduced models.

The ratio 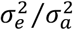 is important when predicting breeding values using the mixed model equations as it is the shrinkage factor for additive effects. The variability of the ratio under different core sizes is shown in Figure 4. As the core size approached eigen98, the ratio became closer to the simulated value of 2.33. Additionally, the ratio became less influenced by the number of generations used to estimate the variance components as the core size approached eigen98. Reliable variance components estimates (or at least their ratio and heritability) are of great importance to ensure the accurate prediction of breeding values. The resulting breeding values are not BLUP unless the true variances are known or are approaching the true parameters (Kennedy, 1981).

**Figure 4.**
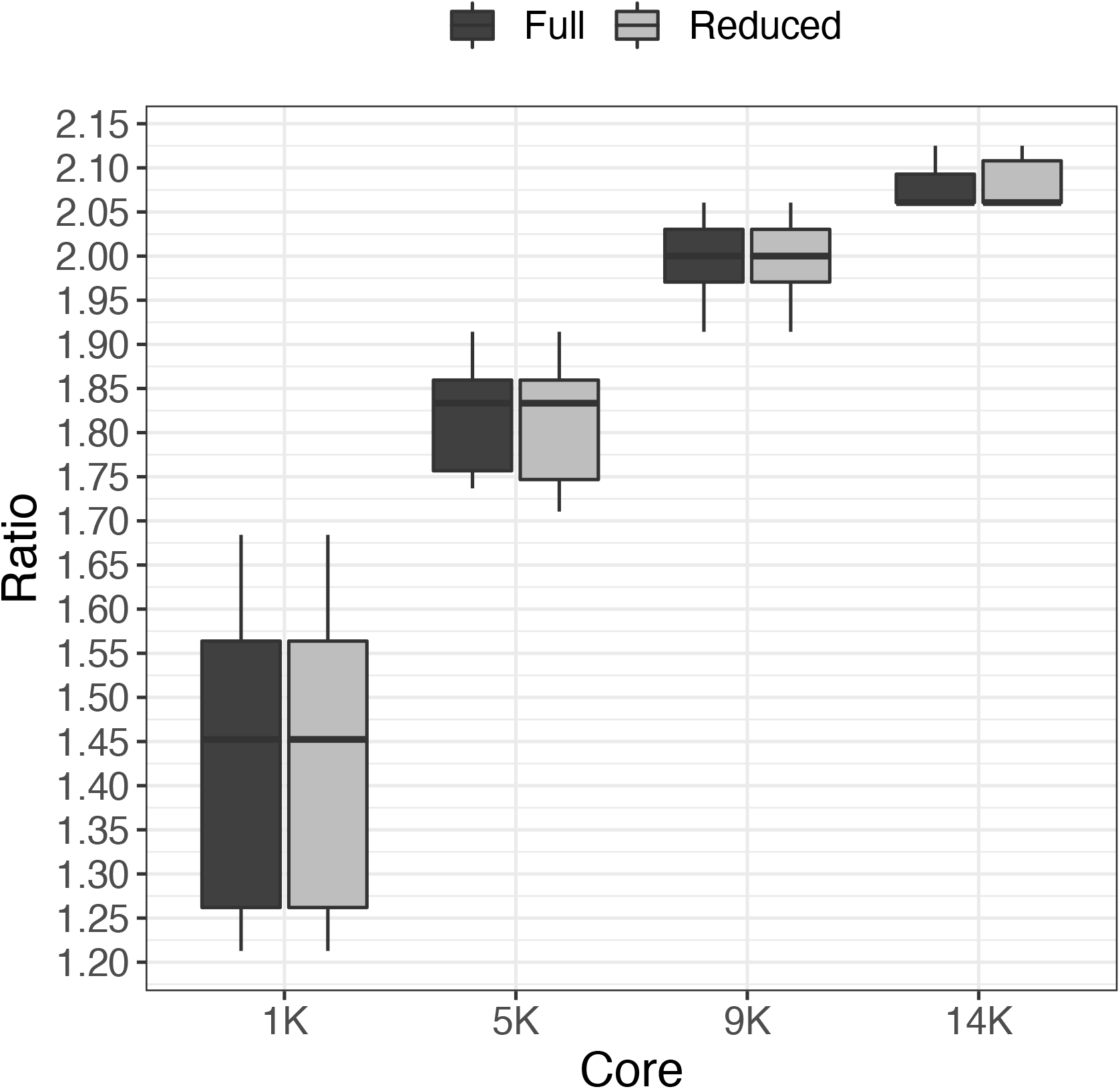
Distribution of the ratio 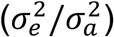 over different number of generations with pedigree and phenotypic data using different sizes for the core group in APY. Two scenarios were considered, where zeros were stored (Full) or not (Reduced). Error bars represents the standard error of prediction under REML.

The adoption of a core group that explains less than eigen98 affected the ability to represent all the independent chromosome segments segregating in the population, traceback gene frequencies, and consequently, accurately establish covariances between genotypic values. In this study, we might have three different sources of changes for genetic variances. The first source is related to the lack of relationships because generations were sequentially removed in different scenarios. Unknown relationships (i.e., incorrect base population definition) affect the estimation of Mendelian sampling variance in different intensities depending on the number of known parents. If both parents are unknown, Mendelian sampling is equal to 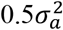, and if only one parent is known, it equals 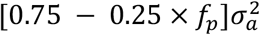, where *f*_*p*_ is the inbreeding coefficient (Henderson, 1976). Under mixed models, offspring breeding values are estimated as a function of parent breeding values and Mendelian sampling. Thus, all individuals with unknown relationships are treated as samples from the base population with average breeding value of 0 and common variance 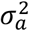.

The second source of change in genetic variance is the presence of selection over generations, which affects the distribution of sire and dam breeding values. Unfortunately, it is impossible to identify the contribution of each factor separately because this study was not designed for that purpose. The third source of genetic variation, which is the aim of this study, is the intentional use of a sparse representation of **G**^−**1**^, i.e., 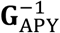. In APY, it is intrinsically assumed that the complete genome is divided into many independent chromosome segments (ICS) containing non-redundant genomic information. The number of ICS is a statistical concept that depends on the effective population size and the genome length (Stam, 1980). The consequence of this assumption is that a small error in variance components estimation can be observed by building the core group considering the dimensionality of **G** as a function of the number of eigenvalues explaining a certain proportion of variance. For example, if 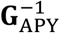 is built based on the number of core animals equal to that of eigenvalues explaining 98% of the variance in **G**, the assumed error is 2% (Misztal et al., 2020). Results from the current study add a new dimension to the factors driving the estimation of reliable variance components in the genomic era. Thus, if the definition of the core group considers the genetic architecture of the population, **G** might contain all the genetic information necessary to estimate reliable variance components (Junqueira et al., 2017; Junqueira et al., 2020). In addition to the factors evaluated in this study, Cesarani et al. (2019) have found that the selection design and genotyping structure can influence bias in estimating variance components.

### Computing resources

Nowadays, much effort has been placed on developing faster and computationally feasible methods for a virtually unlimited number of genotyped individuals. Using large-scale datasets becomes more problematic when the objective is to estimate variance components. This is because most algorithms require several rounds of inversion of the LHS of MME before the convergence is reached. During computations, factorization and inversion are the most demanding steps in the REML estimation. The possibility to combine APY to compute a sparse representation of **G**^−**1**^, data reduction, and YAMS (i.e., dense blocks operation) (Masuda et al., 2014; Masuda et al., 2015) seems computationally beneficial. In this study, we evaluated the factors impacting the timing required for computational operations. Figure 5 shows the average computing time, relative to total (i.e., in percentage), required for ordering, factorization (symbolic and numerical), and sparse inversion with data reduction (pedigree and phenotypes). The most time-consuming operation was the inversion, which took more than 50% of the total time. This was expected because matrix inversion has a cubic computing cost. Next, numerical factorization consumed nearly 30% of the total computing time, whereas ordering and symbolic factorization took approximately 9% and 7.5%, respectively. Skipping zero elements in the MME did not improve the computing time of any of the inverse operations.

**Figure 5.**
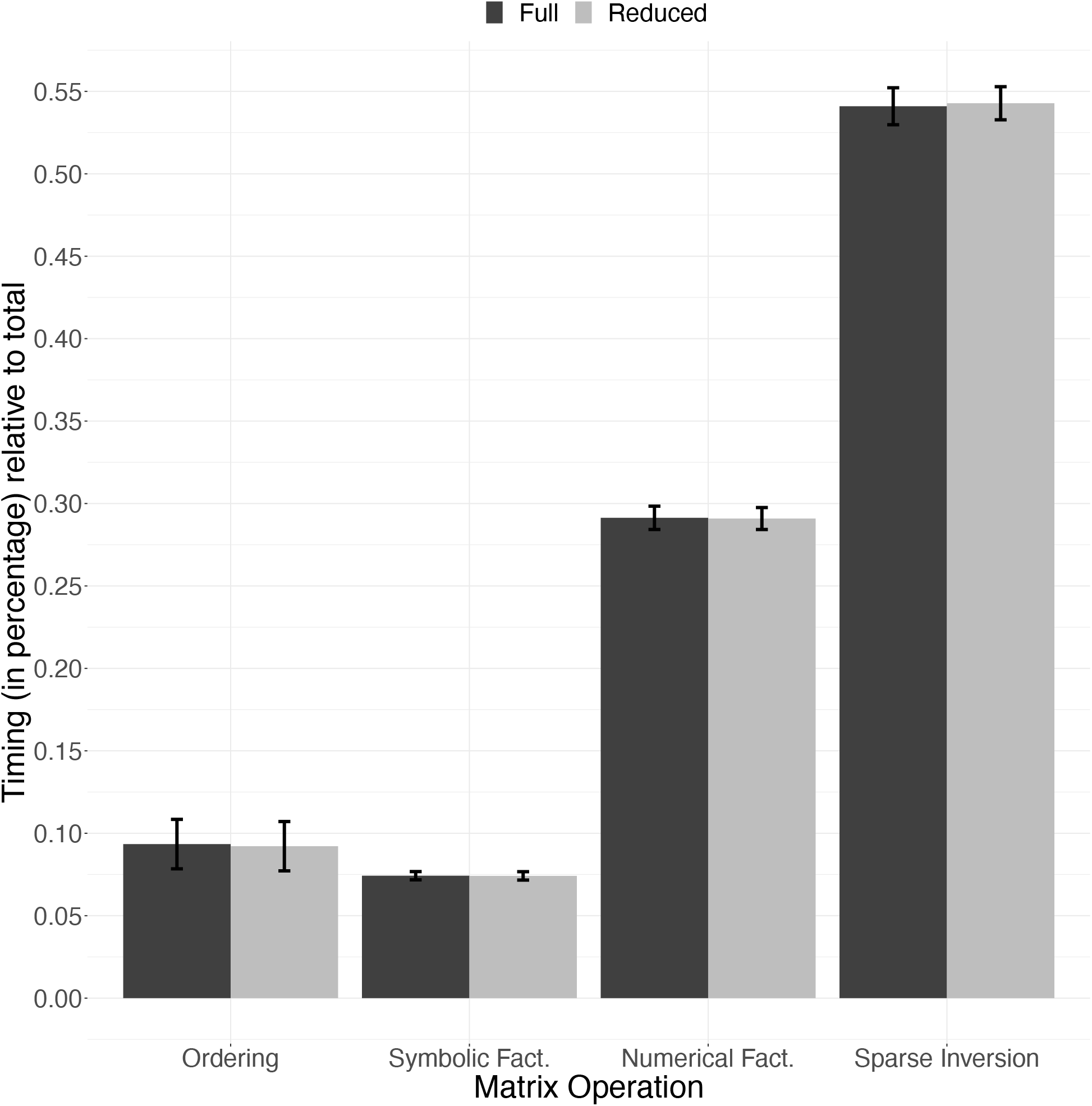
Average timing in percentage (ratio between total timing) relative to each operation used in the process of matrix inversion. The average timing and error bars (standard deviation) were calculated across scenarios using different number of generations in the pedigree and phenotypic and core sizes. The x-axis represents the steps required to invert matrices: finding the ordering, symbolic factorization (Symbolic Fact., setting up the data structure), numerical factorization (Numerical Fact.), and sparse inversion. Two scenarios were considered, where zeros were stored (Full) or not (Reduced).

A detailed description of the computing time required by each step after data removal is in Figure 6. The descriptive statistics of computing time savings across generations is shown in Table 2. Ordering showed the most prominent timing decrease due to data removal, followed by symbolic factorization among the four steps. On average, a 7% decrease in the computing time for ordering was observed by removing each generation of data. During MME computations, ordering and symbolic factorization are not mandatory. These operations are mainly implemented to reduce computing time for numerical factorization and inversion. As more genotypes and/or pedigree records are included in the model, the time required for numerical factorization and sparse inversion increases. Using a simulated dataset with 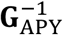 and YAMS, we observed an opposite behavior where shorter pedigree sometimes caused an increase in computing time for the numerical factorization and sparse inversion operations. In these operations, there were no gains in computing performance due to data removal, as shown by the regression slope, which was close to 0 (Table 2). The greatest savings were around 10% when using six generations of pedigree and phenotypic data. It is known that numerical factorization and sparse inversion are the most demanding operations in REML computations. The fact that the required time for these operations was not reduced can be explained by the creation of nonzero elements not present in the coefficient matrix before the numerical factorization is done. Those elements are known as “fill-in elements.”

**Table 2.**
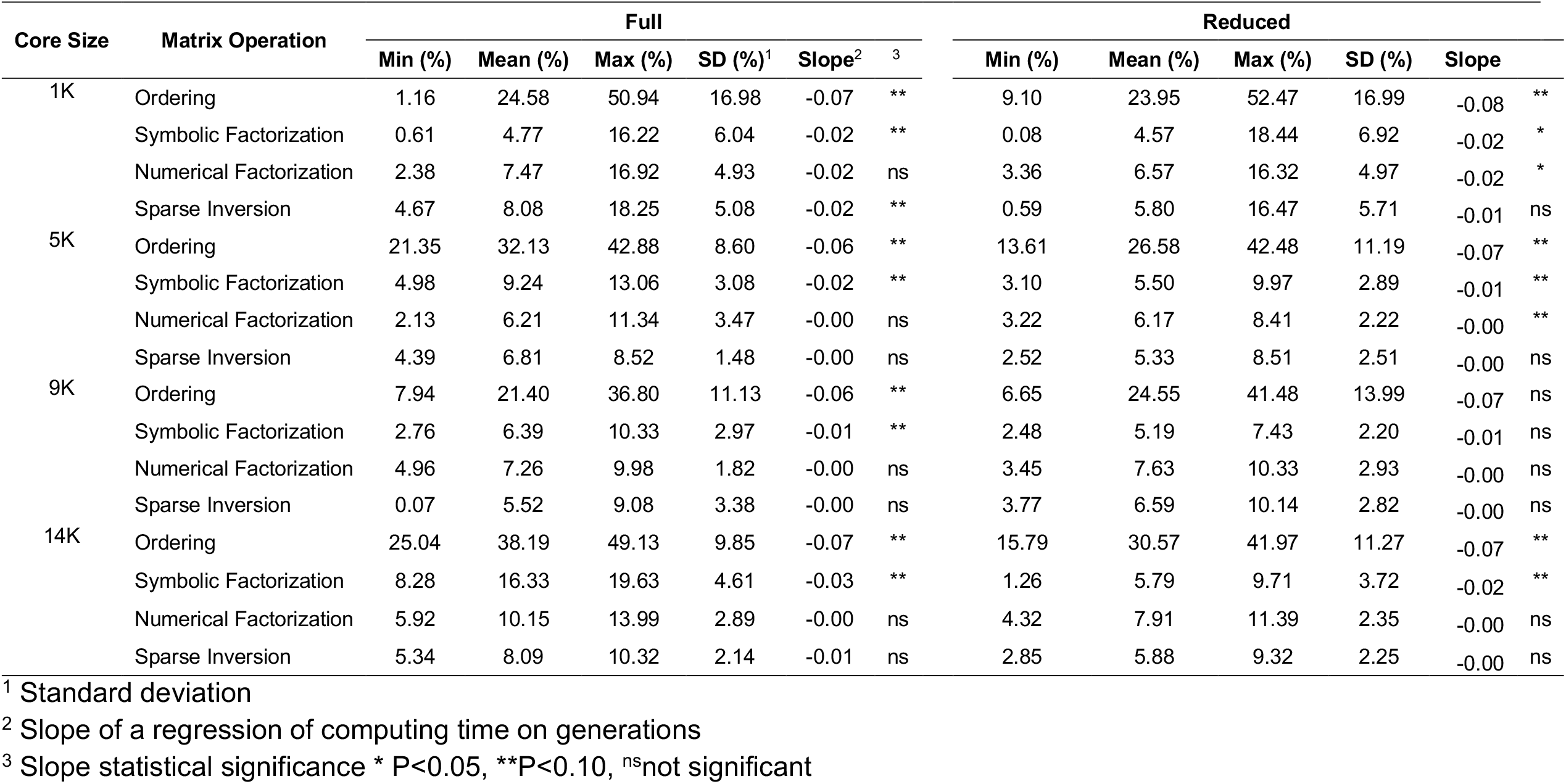
Descriptive statistics of computing time savings for the matrix operations and the slope of a regression of computing time on generations after removing pedigree and phenotypic data. The benchmark is the model using full pedigree and phenotypic data. The comparison is based on using core group of different sizes in algorithm for proven and young (APY), and based on a full mixed model equations (Full) and a reduced mixed model equations after skipping zero elements (Reduced).

**Figure 6.**
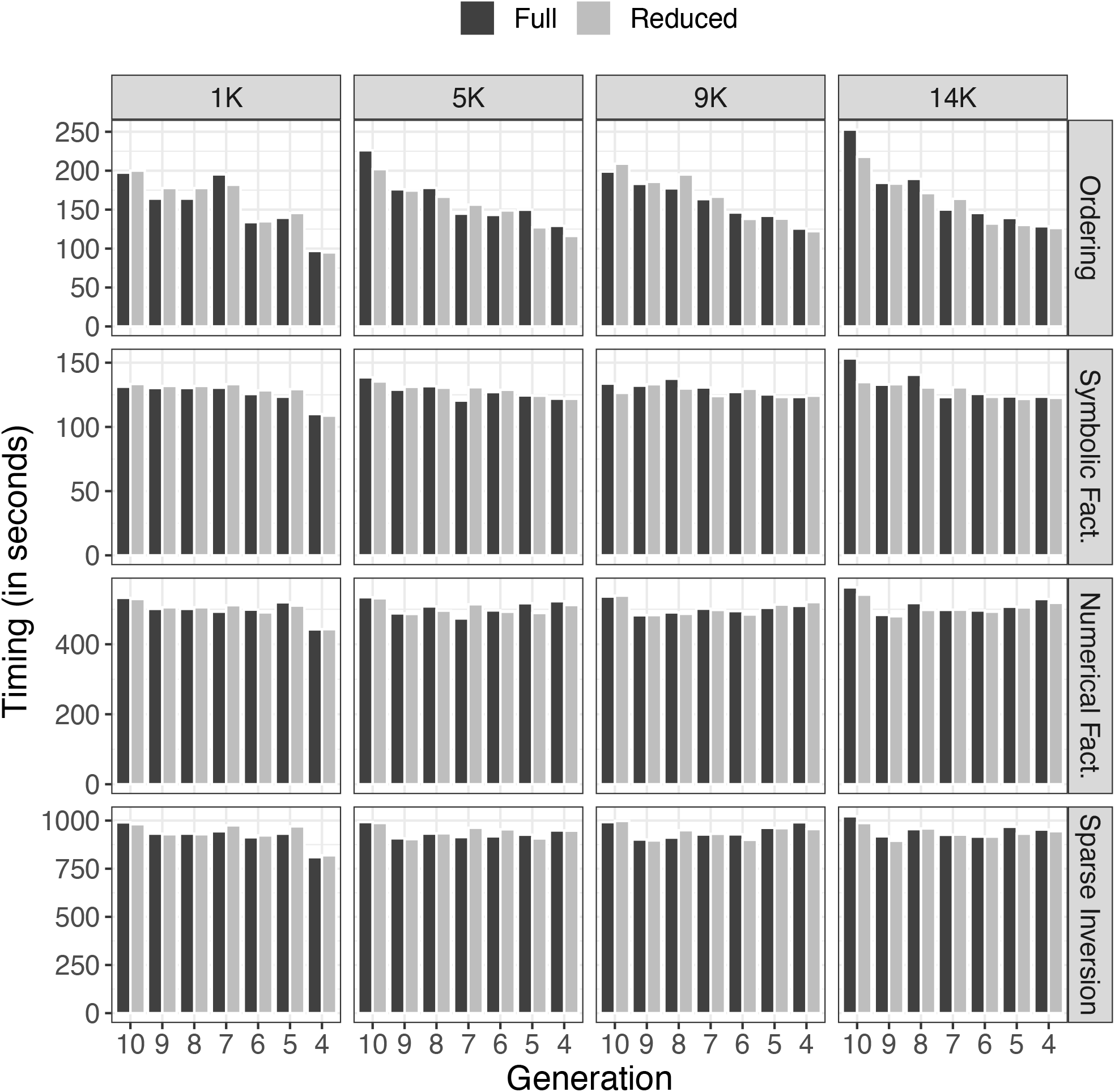
Timing (in seconds) relative to each operation to invert matrices using different number of generations in the pedigree and phenotypes under different number of core animals for the computation of APY **G**^-1^. Matrix inversion steps: finding the ordering (Ordering), symbolic factorization (Symbolic Fact.), numerical factorization (Numerical Fact.), and sparse inversion. Two scenarios were considered, where zeros were stored (Full) or not (Reduced).

Consequently, extra calculations are needed, obviously increasing the amount of time to complete the sparse inversion. There are several efforts in developing faster algorithms focused on typical nonzero structures in sparse matrices. The sparse matrix algorithm in YAMS uses supernodal techniques (i.e., common nonzero pattern between adjacent columns) to speed-up computations. Computing time might be significantly improved compared to other sparse matrix packages (e.g., FSPAK) because the memory hierarchy is more effectively exploited in dense operations, and multiple columns within a submatrix are simultaneously updated (Masuda et al., 2014).

## Conclusions

The algorithm for proven and young (APY) can be successfully applied to create the inverse of the genomic relationship matrix used in single-step genomic restricted maximum likelihood for estimating variance components. To ensure reliable variance component estimation, it is important to use a core size that corresponds to the number of largest eigenvalues explaining around 98% of total variation in **G**. When APY is used, pedigrees can be truncated to increase the sparsity of **H** and slightly reduce computing time for ordering and symbolic factorization, with no impact on the estimates. A reduction in computing time for numerical factorization and sparse inversion is unlike because of fill-in elements. The savings in computing time for estimating variance components is far less than the expected efficiency that APY has shown compared to the use of regular **G**^-1^ for breeding values estimation. This inefficiency is because the block implementation of APY is still not possible for variance components estimation.

## Conflict of interest statement

The authors declare that they do not have any conflict of interest.

## Abbreviations

A: pedigree relationship matrix
AIREML: average information restricted maximum likelihood
APY: algorithm for proven and young
BLUP: best linear unbiased prediction
EBV: estimated breeding value
G: genomic matrix
G_APY_: genomic matrix created using APY
GEBV: genomic enhanced breeding value
GREML: genomic restricted maximum likelihood
IOD: iteration on data
LHS: left hand side of mixed model equations
MME: mixed model equations
QTL: quantitative trait loci
REML: restricted maximum likelihood
ssGBLUP: single step genomic BLUP
ssGREML: single step genomic restricted maximum likelihood
YAMS: yet another MME solver

## Notes

### Competing Interest Statement

The authors have declared no competing interest.

## Literature Cited

Aguilar, I., E. N. Fernandez, A. Blasco, O. Ravagnolo, and A. Legarra. 2020. Effects of ignoring inbreeding in model-based accuracy for BLUP and SSGBLUP. Journal of Animal Breeding and Genetics 137(4):356–364.

Aguilar, I., I. Misztal, D. L. Johnson, A. Legarra, S. Tsuruta, and T. J. Lawlor. 2010. Hot topic: A unified approach to utilize phenotypic, full pedigree, and genomic information for genetic evaluation of Holstein final score. Journal of Dairy Science 93(2):743–752. doi: 10.3168/jds.2009-2730

Bradford, H., I. Pocrnić, B. Fragomeni, D. Lourenco, and I. Misztal. 2017. Selection of core animals in the algorithm for proven and young using a simulation model. Journal of Animal Breeding and Genetics 134(6):545–552.

Cesarani, A., G. Gaspa, F. Correddu, M. Cellesi, C. Dimauro, and N. Macciotta. 2019. Genomic selection of milk fatty acid composition in Sarda dairy sheep: Effect of different phenotypes and relationship matrices on heritability and breeding value accuracy. Journal of Dairy Science 102(4):3189–3203.

Colleau, J. J. 2002. An indirect approach to the extensive calculation of relationship coefficients. Genetics Selection Evolution 34(4):409.

Fragomeni, B. O., D. A. L. Lourenco, S. Tsuruta, Y. Masuda, I. Aguilar, A. Legarra, T. J. Lawlor, and I. Misztal. 2015. Hot topic: Use of genomic recursions in single-step genomic best linear unbiased predictor (BLUP) with a large number of genotypes. Journal of Dairy Science 98(6):4090–4094. doi: 10.3168/jds.2014-9125

Henderson, C. R. 1975. Best linear unbiased estimation and prediction under a selection model. Biometrics:423–447.

Henderson, C. R. 1976. A simple method for computing the inverse of a numerator relationship matrix used in prediction of breeding values. Biometrics:69–83.

Hidalgo, J., D. Lourenco, S. Tsuruta, Y. Masuda, S. Miller, M. Bermann, A. L. Garcia, and I. Misztal. 2021. Changes in genomic predictions when new information is added. Journal of Animal Science 99(2):skab004.

Junqueira, V. S., F. F. Cardoso, M. M. Oliveira, B. P. Sollero, F. F. Silva, and P. S. Lopes. 2017. Use of molecular markers to improve relationship information in the genetic evaluation of beef cattle tick resistance under pedigree-based models. Journal of Animal Breeding and Genetics 134(1):14–26.

Junqueira, V. S., P. S. Lopes, D. Lourenco, F. F. e Silva, and F. F. Cardoso. 2020. Applying the metafounders approach for genomic evaluation in a multibreed beef cattle population. Frontiers in Genetics 11

Kennedy, B. 1981. Variance component estimation and prediction of breeding values. Canadian Journal of Genetics and Cytology 23(4):565–578.

Lidauer, M., I. Strandén, E. A. Mäntysaari, J. Pösö, and A. Kettunen. 1999. Solving large testday models by iteration on data and preconditioned conjugate gradient. J. Dairy Sci. 82(12):2788–2796.

Lourenco, D. A., B. O. Fragomeni, S. Tsuruta, I. Aguilar, B. Zumbach, R. J. Hawken, A. Legarra, and I. Misztal. 2015. Accuracy of estimated breeding values with genomic information on males, females, or both: an example on broiler chicken. Genetics Selection Evolution 47(1):56.

Lourenco, D. A. L., A. Legarra, S. Tsuruta, D. Moser, S. Miller, and I. Misztal. 2018. Tuning indirect predictions based on SNP effects from single-step GBLUP. Interbull Bulletin (53)

Masuda, Y., I. Aguilar, S. Tsuruta, and I. Misztal. 2015. Technical note: Acceleration of sparse operations for average-information REML analyses with supernodal methods and sparsestorage refinements. Journal of Animal Science 93(10):4670–4674.

Masuda, Y., T. Baba, and M. Suzuki. 2014. Application of supernodal sparse factorization and inversion to the estimation of (co) variance components by residual maximum likelihood. Journal of Animal Breeding and Genetics 131(3):227–236.

Masuda, Y., I. Misztal, A. Legarra, S. Tsuruta, D. A. L. Lourenco, B. O. Fragomeni, and I. Aguilar. 2017. Avoiding the direct inversion of the numerator relationship matrix for genotyped animals in single-step genomic best linear unbiased prediction solved with the preconditioned conjugate gradient. Journal of Animal Science 95(1):49–52.

Masuda, Y., I. Misztal, S. Tsuruta, A. Legarra, I. Aguilar, D. A. L. Lourenco, B. O. Fragomeni, and T. J. Lawlor. 2016. Implementation of genomic recursions in single-step genomic best linear unbiased predictor for US Holsteins with a large number of genotyped animals. Journal of Dairy Science 99(3):1968–1974.

Meyer, K. 1997. An average information restricted maximum likelihood algorithm for estimating reduced rank genetic covariance matrices or covariance functions for animal models with equal design matrices. Genetics Selection Evolution 29(2):97.

Misztal, I. 2016. Inexpensive computation of the inverse of the genomic relationship matrix in populations with small effective population size. Genetics 202(2):401–409.

Misztal, I., H. L. Bradford, D. A. L. Lourenco, S. Tsuruta, Y. Masuda, A. Legarra, and T. J. Lawlor. 2017. Studies on inflation of GEBV in single-step GBLUP for type. Interbull Bulletin (51):38–42.

Misztal, I., A. Legarra, and I. Aguilar. 2014. Using recursion to compute the inverse of the genomic relationship matrix. Journal of Dairy Science 97(6):3943–3952.

Misztal, I., S. Tsuruta, I. Pocrnic, and D. Lourenco. 2020. Core-dependent changes in genomic predictions using the algorithm for proven and young in single-step genomic best linear unbiased prediction. Journal of Animal Science 98(12):skaa374.

Misztal, I., S. Tsuruta, T. Strabel, B. Auvray, T. Druet, and D. H. Lee. 2002. BLUPF90 and related programs. In: Proceedings of the 7th World Congress on Genetics Applied to Livestock Production

Patry, C., and V. Ducrocq. 2011. Evidence of biases in genetic evaluations due to genomic preselection in dairy cattle. Journal of Dairy Science 94(2):1011–1020.

Patterson, H. D., and R. Thompson. 1971. Recovery of inter-block information when block sizes are unequal. Biometrika 58(3):545–554.

Pocrnic, I., D. A. Lourenco, Y. Masuda, A. Legarra, and I. Misztal. 2016a. The dimensionality of genomic information and its effect on genomic prediction. Genetics 203(1):573–581.

Pocrnic, I., D. A. L. Lourenco, Y. Masuda, A. Legarra, and I. Misztal. 2016b. The dimensionality of genomic information and its effect on genomic prediction. Genetics 203(1):573–581.

Pocrnic, I., D. A. L. Lourenco, Y. Masuda, and I. Misztal. 2016c. Dimensionality of genomic information and performance of the Algorithm for Proven and Young for different livestock species. Genetics Selection Evolution 48(1):82.

Sargolzaei, M., J. Chesnais, and F. Schenkel. 2011. FImpute-An efficient imputation algorithm for dairy cattle populations. Journal of Dairy Science 94(1):421.

Solberg, T., A. Sonesson, J. Woolliams, and T. Meuwissen. 2008. Genomic selection using different marker types and densities. Journal of Animal Science 86(10):2447–2454.

Stam, P. 1980. The distribution of the fraction of the genome identical by descent in finite random mating populations. Genetics Research 35(2):131–155.

Strandén, I., and E. A. Mäntysaari. 2014. Comparison of some equivalent equations to solve single-step GBLUP. In: Proceedings of the 10th World Congress on genetics applied to Livestock production. Vancouver. p 22.

Tsuruta, S., T. Lawlor, D. Lourenco, and I. Misztal. 2021. Bias in genomic predictions by mating practices for linear type traits in a large-scale genomic evaluation. Journal of Dairy Science 104(1):662–677.

Tsuruta, S., I. Misztal, and I. Stranden. 2001. Use of the preconditioned conjugate gradient algorithm as a generic solver for mixed-model equations in animal breeding applications. J. Anim. Sci. 79(5):1166–1172.

Vandenplas, J., M. P. Calus, and J. Ten Napel. 2018. Sparse single-step genomic BLUP in crossbreeding schemes. Journal of Animal Science 96(6):2060–2073.

VanRaden, P. M. 2008. Efficient methods to compute genomic predictions. Journal of Dairy Science 91(11):4414–4423.

